# Intracranial neural representation of phenomenal and access consciousness in the human brain

**DOI:** 10.1101/2024.04.05.588216

**Authors:** Zepeng Fang, Yuanyuan Dang, Xiaoli Li, Qianchuan Zhao, Mingsha Zhang, Hulin Zhao

## Abstract

After more than 30 years of extensive investigation, impressive progress has been made in identifying the neural correlates of consciousness (NCC). However, the functional role of spatiotemporally distinct consciousness-related neural activity in conscious perception is debated. Based on empirical EEG findings, e.g., of the enhanced early negative wave and late positive wave under conscious conditions, an influential framework proposed that consciousness-related neural activities could be dissociated into two distinct processes: phenomenal and access consciousness. This framework has been supported mainly by comparison of neural activity between report and no-report paradigms; however, though hotly debated, its authenticity has not been examined in a single paradigm with more informative intracranial recordings. In the present study, we employed a novel visual awareness task and recorded the local field potential (LFP) of epilepsy patients with electrodes implanted in cortical and subcortical regions. Overall, we found that the latency of visual awareness-related activity exhibited a bimodal distribution, and the recording sites with short and long latencies were largely separated in location, except in the lateral prefrontal cortex (lPFC). The mixture of short and long latencies in the lPFC indicates that it plays a critical role in linking phenomenal and access consciousness. However, the division between the two is not as simple as the central sulcus, as proposed previously. Moreover, in 4 patients with electrodes implanted in the bilateral prefrontal cortex, early awareness-related activity was confined to the contralateral side, while late awareness-related activity appeared on both sides. Finally, Granger causality analysis showed that awareness-related information flowed from the early sites to the late sites. These results provide the first LFP evidence of neural correlates of phenomenal and access consciousness, which sheds light on the spatiotemporal dynamics of NCC in the human brain.

## Introduction

Identifying the neural correlates of consciousness (NCC) is an intriguing and arduous task for modern neuroscience research^1^. During the past three decades, the neural correlates of visual awareness, a core component of consciousness in primates, have been studied extensively^2-4^. Functional magnetic resonance imaging (fMRI) studies have found that the frontoparietal network^5^, temporal lobe^6^, insula and anterior cingulate gyrus^7,8^ are associated with visual awareness. In addition, electroencephalography (EEG) studies^9-12^ have identified two components of event-related potentials (ERPs) that are highly correlated with visual awareness: the early visual awareness negativity (VAN), which appears ∼200 ms after visual stimulus onset, and the late P3b, which appears ∼350 ms after visual stimulus onset. Furthermore, a few intracranial recording studies in patients with implanted electrodes have also revealed the neural correlates of visual awareness in multiple brain regions; for example, awareness-related activity starts at approximately 200 ms and 300 ms in the medial temporal lobe^13,14^ and prefrontal cortex^15^, respectively.

While the findings of these studies provide valuable evidence of NCC, the functional role of spatiotemporally distinct awareness-related activity in conscious awareness is hotly debated^16,17^. This controversy centers on differences in definitions of NCC, i.e., the minimal neuronal mechanisms jointly sufficient for any one specific conscious experience. Some investigators have argued that perceptual content must be accessible to a wide range of cognitive processes to form specific conscious experiences; therefore, NCC should contain consciousness-related cognitive activity^1,12^. In contrast, other investigators have questioned the necessity of cognitive access in consciousness and argued that cognitive access is the consequence of consciousness but not the conscious experience per se^16,17^.

A recent framework seems to provide a promising solution for the above controversy by dividing consciousness into phenomenal consciousness (p-consciousness) and access consciousness (a-consciousness)^18^. This framework involves two distinct components of consciousness: p-consciousness, which refers to subjective experience (so-called ‘‘qualia’’) without further cognitive processing, and a-consciousness, which refers to the consequences of subjective experience, such as working memory, decision-making and motor planning. The most powerful evidence that supports this framework was provided by studies that compared NCC between report (requiring cognitive processes) and no-report (not requiring cognitive processes) paradigms. In EEG studies, the late-appearing NCC (e.g., P3b), which peaks in anterior brain areas, was absent in no-report paradigms but not in report paradigms; however, the early-appearing NCC (e.g., VAN), which peaks in posterior brain areas, was consistent between no-report and report paradigms^19^. Consistent with these findings, in fMRI studies, consciousness-related activity in the prefrontal cortex was largely weaker in no-report paradigms than in report paradigms, whereas consciousness-related activity in posterior brain areas was similar between the two types of paradigms^20^. Therefore, some investigators have argued that early consciousness-related activity in posterior brain areas is involved in p-consciousness and that late activity in the prefrontal cortex is involved in a-consciousness^21,22^. However, the importance and authenticity of this framework has been doubted, i.e. some researchers argued that there is only one, but not two, NCC^23-25^. Moreover, it is obvious that limiting comparisons to report and no-report paradigms restricted the exploration of the spatiotemporal dynamics of NCC between p- and a-consciousness. In addition, the frequently used noninvasive techniques (EEG, fMRI, etc.) cannot precisely map the spatiotemporal dynamics of NCC due to limitations in spatial or temporal resolution.

To address these questions, we recorded the LFP in 9 epilepsy patients by implanting sEEG electrodes (1334 recording sites) across multiple brain regions, including cortical and subcortical areas, while they performed a novel report-based visual awareness task (Figure 1). We thus explored the spatiotemporal dynamics of visual consciousness (awareness)-related activities between p- and a-consciousness among multiple brain regions.

**Figure 1.**
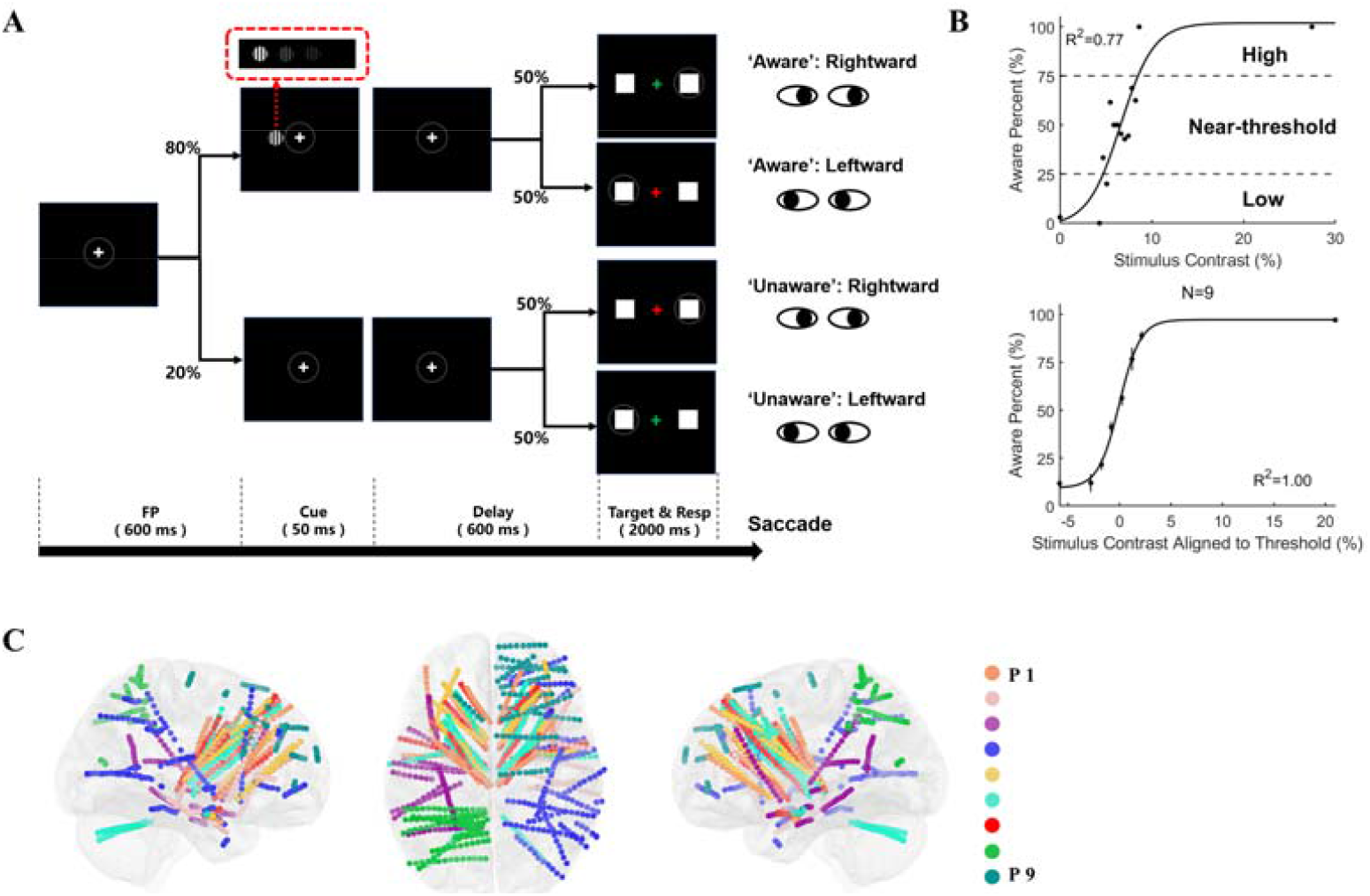
Visual awareness task: Behavioral results, and electrode localization A. Schematic of the visual awareness task. A trial started when a fixation point (0.5° × 0.5°, white cross) appeared at the center of the screen (radius of eye position check window = 4°, shown as the dotted circle). After the subject fixated on the fixation point for 600 ms, a cue stimulus (Gabor grating, 2 × 2° circle) was presented for 50 ms at a fixed position (7°) on the left (or right, see Methods) side of the screen. In 70% of the trials, the grating contrast was maintained near the subject’s perceptual threshold by a staircase method; in 10% of the trials, the stimulus contrast was well above the threshold; and in the other 20% of the trials, the stimulus contrast was 0, namely, no stimulus appeared. After another 600 ms delay, the color of the fixation point turned red or green, and two saccade targets (1 × 1°, white square) appeared at fixed positions (10°) on the left and right sides of the screen. If the grating was seen, a green fixation point indicated that subjects should make a saccade to the right target, while a red fixation point suggested that subjects should make a saccade to the left target. If the grating was not seen, the rule of saccadic direction was inverted. B. Psychometric detection curves; the upper panel shows an example session of a patient, and the lower panel shows the population data of all patients. Each black point in the graph represents the awareness percentage in a correlated contrast level, and the black curve represents the fitted psychometric function. Awareness percentages greater than 25% and less than 75% are defined as near-threshold, whereas awareness percentages less than 25% are defined as low and those greater than 75% are defined as high. In the lower panel, the contrast is aligned to the individual subject’s perceptual threshold (50% awareness percentage), i.e., the contrast 0 represents each subject’s perceptual threshold. N, number of patients; R^2^, coefficient of determination. C. Left, top and right views of all recording sites projected on an MNI brain template. Each color represents a patient. In all brain images, the right and upper sides of the image represent the right and upper sides of the brain.

## Results

### Behavioral Results

Figure 1B shows the behavioral performance of a representative session from one patient (upper panel) and the population result from all patients (lower panel). The results showed a classic psychometric curve, i.e., as the grating contrast increased, the proportion of individuals reporting that they were ‘aware’ of the grating gradually increased. In trials with no grating or high grating contrast, patients showed high accuracy (94.75% ± 2.65), indicating that the patients understood and performed the task according to the task requirements. In trials with near-threshold contrast, where the grating contrasts were similar, patients varied in reports of being ‘aware’ or ‘unaware’. Such results indicate that the grating contrast level directly impacted the emergence of visual awareness and thus is a reliable way to study visual awareness.

We divided the trials into 4 conditions for further analysis according to the level of grating contrast and the awareness state reported by the patients: the high contrast-aware (HA), near threshold-aware (NA), near threshold-unaware (NU), and low contrast-unaware (LU) conditions (Fig. 2A, see the Methods for details).

**Figure 2.**
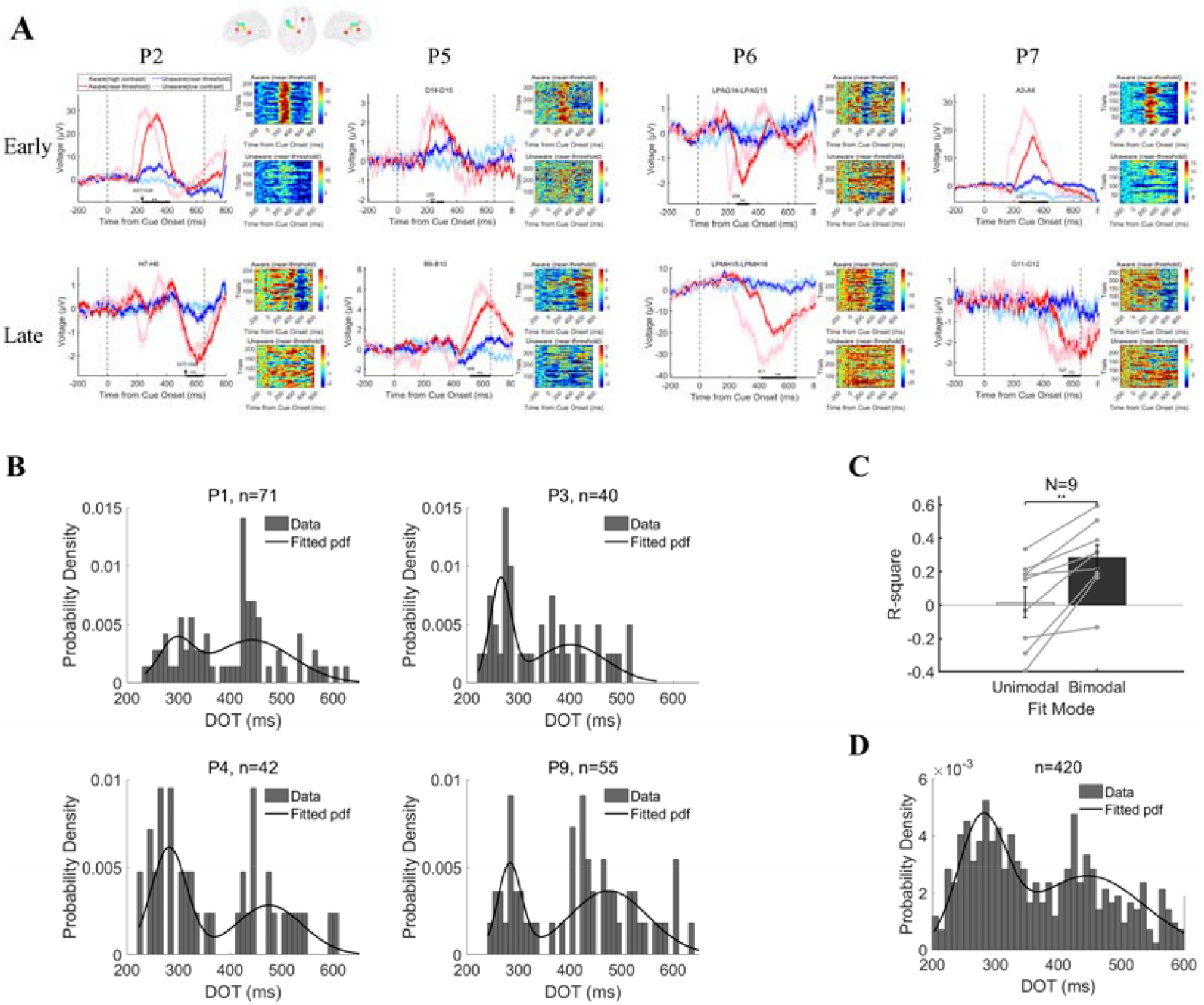
Examples of visual awareness-related activity and the distributions of DOTs. A. Example LFP from 4 patients. In each panel, the left figure shows the grand average of LFP activity in 4 different conditions. The pink/red/blue/light blue lines represent data under the HA/NA/NU/LU conditions, respectively. The shaded area of the curve represents the standard error of the mean (SEM). The two black dashed lines at 0 ms and 650 ms represent the grating onset and fixation point color change (the appearance of saccade targets), respectively. The thick black line represents a significant difference between the NA and NU conditions (p<0.01 corrected, independent sample t test). The right part of each panel shows the LFP amplitude in a single trial under the NA (upper) and NU (lower) conditions. The color bar in each panel represents the LFP amplitude. B. Histogram of the DOT distribution in example patients. The bar plots in the histogram represent the probability density of DOTs in different time bins (10 ms for each time bin), and the black curve represents the result of bimodal fitting. C. Comparison between unimodal and bimodal fitting results. White bars represent the values of R^2^ from unimodal fitting, and black bars represent the values of R^2^ from bimodal fitting. The gray points in the panel represent the R^2^ obtained from the individual patient. The R^2^ values from the same patient are connected by a gray line. R^2^, goodness-of-fit index. D. Histogram of the populational DOT distribution. The bar plots in the histogram represent the probability density of DOTs in different time bins (10 ms for each time bin), and the black curve represents the results of bimodal fitting.

### iEEG results

While patients (N = 9) performed the visual awareness task, we recorded the LFP in multiple brain regions (total=1334 recording sites, 420 sites were awareness related), including frontal, parietal, temporal, occipital, and subcortical regions (such as the thalamus). Figure 1D shows the locations of all electrodes in all patients (projected on the Montreal Neurological Institute (MNI) brain template ICBM152).

### The LFP indicates two temporally distinct awareness-related activities

Figure 2A shows representative ERP activity, which reflects the typical ERP activity of 9 patients. The recording sites are shown in different colors in the brain template insert (each color represents the recording sites of an individual patient). The grand average (left) and single-trial (right) ERP data had several notable features. First, early and intense ERP activity was evoked in the HA condition (light red line) compared to the other three conditions (LU, light blue line; NA, red line; NU, blue line). Second and more importantly, in the near-threshold conditions, although the contrasts of the grating were similar between NA and NU trials (as reported previously^26^), the ERP activity in the NA and NU trials started to diverge significantly ∼200 ms after grating onset (the thick black line in the panels denotes p < 0.01, corrected, independent t test; for details see the Methods). The ERP activity difference between NA and NU trials was robust and constant across all 9 patients, as demonstrated in single trial analysis (the right side of Figure 3A shows data from more than 100 trials for the NA and NU conditions in individual patients; for details, see the Methods). Because the grating contrast was very similar between NA and NU trials, differences in ERP activity between NA and NU conditions was considered to reflect awareness-related activity. Therefore, the divergence onset time (DOT) represents the latency of visual awareness-related activity at a specific recording site.

**Figure 3.**
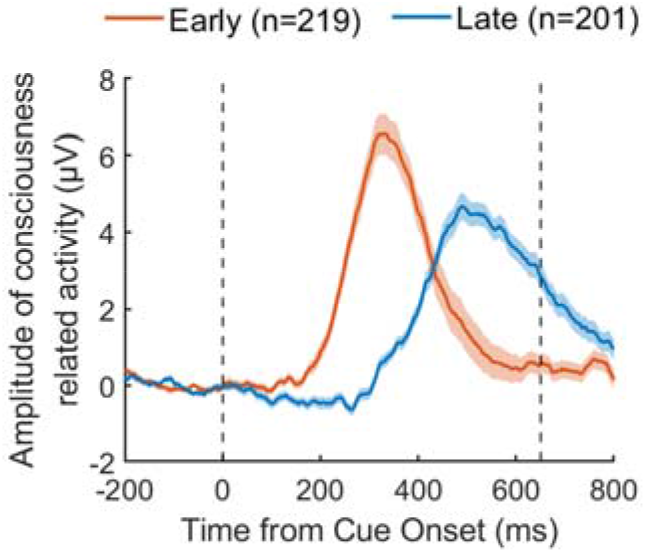
Early and late awareness-related activities in the population. The orange line represents early awareness-related activity, and the blue line represents late awareness-related activity. The shaded area around each line represents the standard error of the mean (SEM). The two black dashed lines at 0 ms and 650 ms represent the grating onset and fixation point color change onset (the appearance of saccade targets), respectively. N, number of recording sites.

Notably, in Figure 2A, the DOTs in upper panels are shorter than in lower panels in each individual patient. To further characterize the temporal features of DOTs, we first calculated the distribution of DOTs for individual patients in all recording sites with visual awareness-related activity (Fig. 2B). The DOTs in the example patients tended to show a bimodal distribution rather than a unimodal distribution. To quantify this tendency, we fitted the distribution with unimodal and bimodal functions. The results showed that the coefficient of determination (R^2^) of the bimodal fitting function was significantly greater than that of the unimodal fitting function in all 9 patients (p < 0.01, Fig. 2C). Such results indicate that the distribution of DOTs was better characterized by a bimodal distribution than by a unimodal distribution in individual patients. Then, we pooled the DOTs of all patients and calculated the DOT distribution of all visual awareness-related sites (n = 420). The distribution of DOTs also showed a significant bimodal distribution (p < 0.01); DOTs divided into early (n = 219) and late (n = 201) activity according to the cutoff value (t = 368 ms) (Figure 2D).

To further assess the temporal profile of awareness-related activity, we calculated the ERP activity of early and late DOTs in the population of 9 patients. Interestingly, early awareness-related activity ceased at 400-500 ms after grating onset, which is well before the next stage of the task (the time that the fixation point changed color). Such results indicate that early awareness-related activity is not directly involved in subsequent report-related processes. In contrast, late awareness-related activity occurred 650 ms after grating onset, i.e., after the time that the fixation point changed color, and thus may be involved in the postperceptual cognitive process.

### LFP data indicate two spatially distinct awareness-related activities

Next, we assessed the spatial locations of early and late awareness-related activity. Due to the sparse sampling sites of iEEG recording in individual patients and recording site differences among patients, we included only 14 brain regions with sufficient data (see the Methods for details). Then, we analyzed the proportions of early and late sites within each of these 14 regions. To clearly display the locations of early and late awareness-related activity in the brain, their proportions were projected on the brain template. The results are shown in Figure 4A and 4B. As shown in the figure, the locations of early and late awareness-related activity were largely separated, i.e., the early sites were mainly concentrated in the lateral prefrontal cortex (including the pars opercularis and middle frontal gyrus), posterior parietal cortex (including the superior and inferior parietal lobule), thalamus and caudate nucleus; the late sites were mainly concentrated in the caudate nucleus, insula, superior frontal gyrus, and caudal anterior cingulate gyrus. However, the lateral PFC (lPFC) showed similar proportions of early and late sites, which indicates that the lPFC is a node linking early and late awareness-related processes.

**Figure 4.**
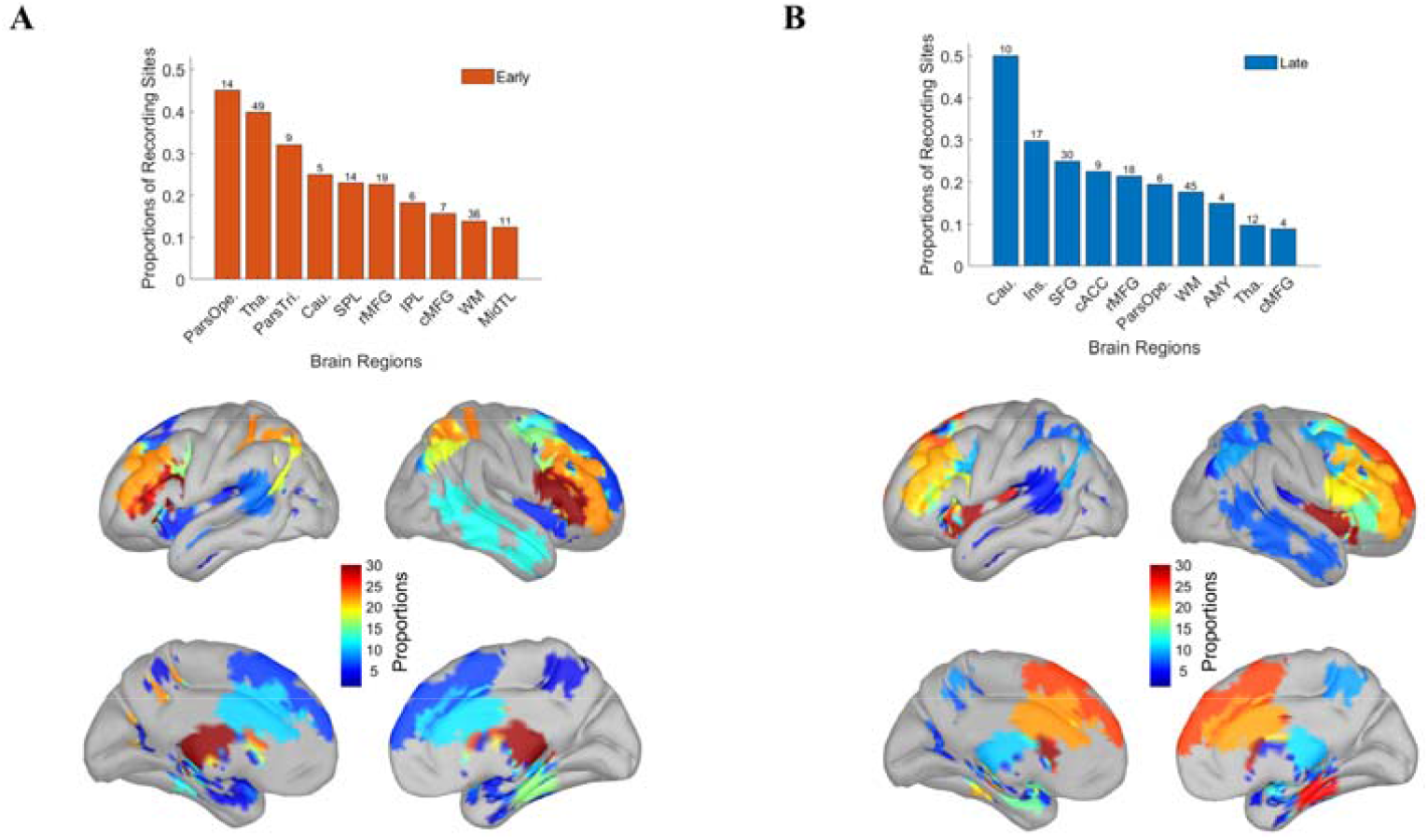
Locations of early and late awareness-related activity A. The proportions of the early awareness-related activities in multiple brain regions. The histogram shows the proportions of early awareness-related activities in different brain regions, sorted by the values of proportion. The number at the top of each bar represents the number of early sites. The topographic map in the lower part of the panel shows the same data as in the histogram. The 4 maps from upper left to lower right represent the left, right, left medial and right medial views, respectively. The color bar represents the value of proportion. Par Ope., pars opercularis; Tha., thalamus; ParsTri., pars triangularis; Cau., caudate; SPL, superior parietal lobule; MidTL, middle temporal gyrus. B. The proportions of the late awareness-related activities in multiple brain regions. Same as Panel A, but for late awareness-related activity. Cau., caudate; Ins., insula; SFG, superior frontal gyrus; cACC, caudal anterior cingulate; rMFG, rostral middle frontal gyrus; ParsOpe., pars opercularis; WM, white matter; AMY, amygdala; Tha., thalamus; cMFG, caudal middle frontal gyrus.

In addition, 4 patients in our study had electrodes implanted in the bilateral PFC, with similar numbers and locations on each side. We further conducted an analysis to assess whether there was a biased distribution of awareness-related activity between the two sides of the PFC. Figure 5 shows that the percentage of early awareness-related activity among recording sites was markedly higher in the contralateral PFC (related to the location of grating) than in the ipsilateral PFC, whereas the percentage of late awareness-related activity was similar between the two sides of the PFC. Such results suggest that early awareness-related activity is mainly confined to the contralateral PFC, whereas late awareness-related activity appears on both sides of the PFC.

**Figure 5.**
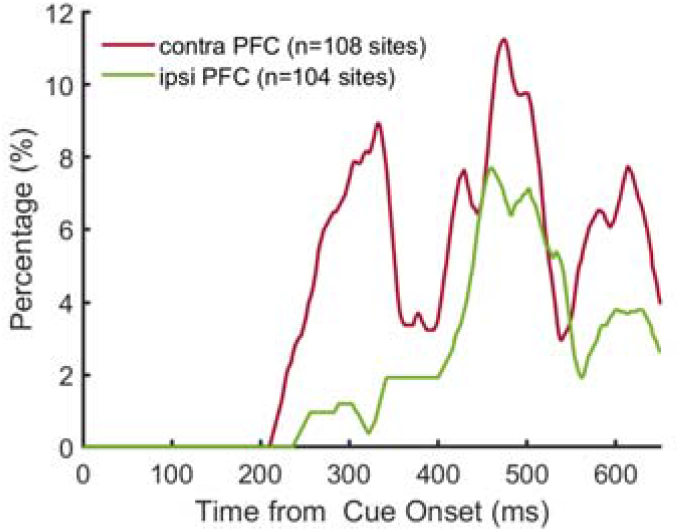
Lateralization of awareness-related activity. The percentages of awareness-related activity in the contralateral and ipsilateral PFC. The red line represents the percentage of awareness-related activity in the contralateral PFC, and the green line represents the percentage of awareness-related activity in the ipsilateral PFC. n, number of recording sites.

### The correlation within and between early and late awareness-related activity

In a previous work^27^, we showed that visual awareness is associated with an increase in the phase-locking value (PLV) in the low-frequency band (1-8 Hz) of the LFP across multiple brain regions, but not in other frequency bands. To assess whether there are correlations within and between early and late awareness-related activity in the present study, we calculated the awareness-related PLV gain in 1-8 Hz, i.e., the PLV difference between the NA and NU conditions, for each paired recording site as a function of time. The data from individual patients showed that the PLV gains started to increase at ∼230 ms after grating onset and had the highest values between early and early activities (Figure 6A, the horizontal and vertical lines divide the early and late activities based on the DOT distribution shown in Figure 2). Then, the PLV gains gradually increased between early and late activities (∼310 ms after grating onset) and finally increased between late and late activities (∼390 ms).

**Figure 6.**
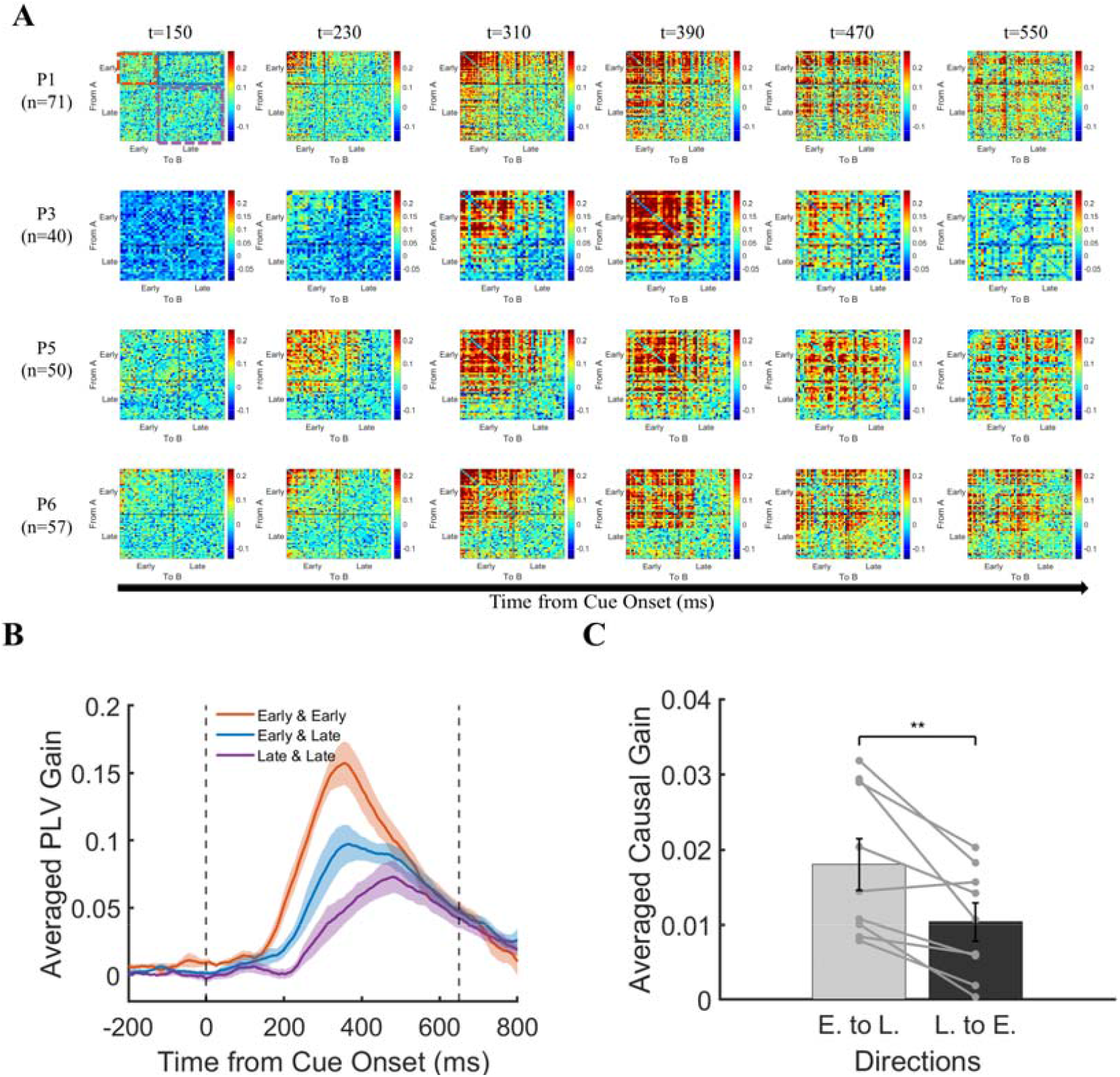
Functional connectivity and information flow between early and late activity A. Temporal dynamics of awareness-related PLV gains in example patients. Heatmaps represent the PLV gains between each pair of LFP signals (sorted by the DOT values, from early to late). The horizontal and vertical gray lines in the heatmap divide early and late awareness-related activity based on the DOT distribution in Figure 2. The dashed colored squares in the upper-left heatmap represent the three PLV conditions: early & early (orange), early & late (blue), and late & late (purple). The heatmaps in each row represent the PLV results from one example patient (P1/P3/P5/P6), and the heatmaps in each column represent the PLV results in a time bin (1 ms) after the onset of the grating (t = 150/230/310/390/470/550 ms). B. Average PLV gains within and between early and late awareness-related activity. The orange/blue/purple lines represent the average PLV gains between early & early, early & late, and late & late awareness-related activity, respectively. The shaded area around each curve represents the SEM. The two black dashed lines at 0 ms and 650 ms represent the grating onset and fixation point color change onset, respectively. C. Average Granger causality gains between early and late activity. The gray bar represents the average GC gains from early to late activity, and the black bar represents the average GC gains from late to early activity. Gray points represent the GC gains of individual patients. The GC gains from the same patient are connected by a gray line. E., early awareness-related activities; L., late awareness-related activities. **, p < 0.01.

To analyze the PLV gain across the population, we first averaged the PLV gain under three conditions, i.e., the early and early, the early and late, and the late and late (indicated by colored squares in Figure 6A), of individual patients at each time point (step width of 1 ms), from -200 to 800 ms of grating onset. Then, we pooled the averaged PLV gains of the 9 patients. The mean (and SEM) PLV gain are shown in Figure 6B. The population results were consistent with the results of individual patients (Figure 6A). The PLV gain between early and early activities increased soon (∼230 ms) after grating onset and had the highest values. The PLV gain between early and late activities started to increase at ∼310 ms and had lower values than those between early and early activities. Finally, the PLV gain between late and late activities started to increase at ∼390 ms and had the lowest values. Such results indicate that early and late awareness-related activities are organized in different functional networks.

### The direction of information flow between early and late awareness-related activities

To further assess the direction of awareness-related information flow between early and late awareness-related activities, we conducted spectral Granger causality (GC) analysis (with a 1-8 Hz band). For each pair of early and late activities, we calculated the GC during 150-650 ms after grating onset (see the Methods for details). Then, we further calculated the GC difference between the NA and NU conditions to determine the awareness-related GC gain. The results of GC gain for each patient in the early-to-late and late-to-early directions are shown in Figure 6C. It is obvious that the GC gains in the early-to-late direction were significantly larger than those in the late-to-early direction. Such results indicate that awareness-related information primarily flows from early awareness-related activity to late awareness-related activity.

## Discussion

### Intracranial LFP activity indicates the presence of two temporally distinct NCC

The present study is the first to report that the latencies of consciousness (awareness)-related activities in multiple brain regions exhibit a bimodal distribution, with modes approximately 200-350 ms (early) and 350-600 ms (late) after grating onset (Figure 2A-D). Furthermore, the early activity ended at 400-500 ms, which is well before the next stage of the task (650 ms after grating onset). Such results indicate that early activity is not directly involved in subsequent report-related processes. In contrast, late activity exceeded the next stage of the task (the time that the fixation point changed color); thus, it may be involved in subsequent report-related processes, such as postperceptual cognitive processes (Figure 3). Therefore, activity at these two times may represent two different consciousness-related processes, i.e., p-consciousness and a-consciousness.

In previous scalp EEG studies, which had high temporal resolution and similar biological origins of the LFP^28^, two ERP components, i.e., the early-occurring VAN and late-occurring P3b, have been considered the most promising NCC candidates^29,30^. The VAN often begins 200-300 ms after stimulus onset (which may be delayed with lower contrast stimuli^31^) and is considered the most consistent neural correlate of p-consciousness^30^. The P3b usually occurs at approximately 350-500 ms (which can vary according to task demands) and is considered a reliable biomarker of a-consciousness^1,12^. However, the aforementioned two assumptions are mainly supported by the comparison between report and no-report paradigms^19^. It is not clear whether there are two distinct activities in the human brain that represent p- and a-consciousness. In the present study, the bimodal distribution of early and late consciousness-related activities and their distinct ending times supported the existence of two different consciousness-related processes in the human brain, i.e., p- and a-consciousness. In addition, the time courses of early and late consciousness-related activity in our study are consistent with the two NCC candidates, i.e., the VAN and P3b. Such similarity across studies suggests that early and late activity based on LFP data might share similar consciousness-related processes with the VAN and P3b, respectively.

### The intracranial LFP activity indicates two NCC largely separated in location

In previous EEG studies, the VAN amplitude has usually peaked in the occipital-temporal areas contralateral to the visual stimulus, while the P3b amplitude has peaked in the more anterior and central frontoparietal areas^10,11^. Although these findings suggest that the VAN and P3b likely originate in different brain regions, we were not able to map the sources of these two components precisely due to the low spatial resolution of scalp EEG. On the other hand, in fMRI studies, the neural correlates of p- and a-consciousness have been mainly distinguished by the comparison between report and no-report paradigms. In report paradigms (including p- and a-consciousness), awareness-related activity has appeared in multiple brain regions, such as the frontoparietal network, middle temporal lobe, insula and temporoparietal junction^6,8,32^. In contrast, in the no-report paradigm (without a-consciousness), fewer brain regions, such as the lateral occipital complex and the middle and inferior frontal gyri, have been activated and are considered to be specifically associated with p-consciousness^8,20,32^. However, the poor temporal resolution of fMRI restricted explorations of the temporal relationship between the processes of p- and a-consciousness.

In the present study, the results show that early and late awareness-related activity was largely separated in location, except in the lateral prefrontal regions that we discuss further in the next section. The locations of early activity were distributed across posterior and anterior regions of the brain, including the middle and inferior frontal gyrus, superior and inferior parietal lobule, and thalamus. These results are largely consistent with findings of previous fMRI studies using no-report paradigms. In addition, the locations of late activity were in more anterior regions of the brain, including the superior and middle frontal gyrus, anterior cingulate, and anterior insula. These results are consistent with findings of report-related activity in previous fMRI studies^8^.

The different anatomical profiles of early and late consciousness-related activity provides further evidence in support of the framework of p- and a-consciousness. Furthermore, although late activity was concentrated in the anterior regions of the brain compared to early activity, the locations exhibiting both early and late activity were not simply divided by the previously proposed ‘front and back’ dichotomy of the brain^29,33^. Rather, it is more likely that these locations are divided according to the different brain-wide networks.

### The intracranial LFP activity suggests that the lPFC is a key node linking p- and a-consciousness

While early (representing p-consciousness) and late (representing a-consciousness) activity is largely separated in location, the lPFC is an exception that is similarly involved in both early and late activity (Figure 4A-B). Such results indicate that the lPFC participates in both p- and a-consciousness. Thus, the lPFC might be a key node in initiating the transformation from p-to a-consciousness. The involvement of the lPFC in a-consciousness has been reported by various studies, including single neuron recording^34-36^, fMRI^5,7^, iEEG^15^ and TMS techniques^37^. However, only a few recent studies have reported the involvement of the lPFC in p-consciousness^8,32,38^. To the best of our knowledge, the involvement of the lPFC in both p- and a-consciousness and its potential role in transforming p-to a-consciousness has not been reported thus far.

### The dynamic relationship between early and late awareness-related activity

We have discussed the temporal and spatial distinctions of the two consciousness-related activities. An interesting question is how early and late activities interact with each other. To answer this question, we conducted an additional analysis. There were 4 patients with electrodes implanted in the bilateral PFC, which allowed us to directly compare the two consciousness-related activities between the hemispheres. We found that early consciousness-related activity was largely confined to the contralateral PFC, whereas late activity appeared on both sides of the PFC (Figure 5A & 5B). Hemispheric differences in consciousness-related activity have been reported by previous scalp EEG studies^10,11^, i.e., the early VAN was found to peak in the contralateral hemisphere, and the late P3b peaked in more central and anterior areas in both hemispheres. Such results suggest that p-consciousness emerges primarily in the contralateral hemisphere, and then consciousness-related information is ‘broadcast’ (a-consciousness) to both hemispheres. Hemispheric differences in visual processing have been reported previously^39^. In this previous study, damage to the lPFC reduced neuronal activity in the extrastriate cortex of the lesioned hemisphere, and the detection ability in the contralesional hemifield was impaired. Although these authors did not explicitly link their results to visual consciousness, the employed paradigm required conscious visual discrimination between targets and distractors. Thus, the results of this study and our study provide cross-validation regarding the hemispheric differences in conscious perception.

Second, the functional connectivity analysis showed that the phase-locking value (PLV) was higher within pairs of early activities but was lower for pairs of early and late activities. Such results indicate that early and late activities involve different functional networks. Moreover, the PLV within pairs of late activities was the lowest. One possible explanation is that late activity is widely distributed among functionally different neural networks, which use p-consciousness information to carry out diverse report-related processes^1^. Finally, the Granger causality analysis showed that consciousness-related information flowed from early sites to late sites.

In summary, our study is the first to provide intracranial LFP evidence of the coexistence of two spatiotemporally distinct consciousness-related activities in multiple brain regions, providing powerful evidence supporting the framework of p- and a-consciousness. Furthermore, our results revealed the dynamic process by which consciousness-related information is transmitted from early to late locations to achieve p- and a-consciousness.

## Methods

### Data acquisition

Nine adult patients (8 males and 1 female, mean age ± SEM: 30.00 ± 5.17 years) with drug-resistant epilepsy or headache participated in this study. All patients had normal or corrected-to-normal vision. Electrophysiological signals (LFPs) were obtained from 9 patients. Stereotaxic EEG depth electrodes (Sinovation Medical Technology Co., Ltd., Beijing, China) containing 8 to 20 sites were implanted in patients at the Department of Neurosurgery, Chinese PLA General Hospital. Each site was 0.8 mm in diameter and 2 mm in length, with 1.5 mm between adjacent sites. A few electrodes were segmented, and the distance between the two segments was 10 mm. Electrode placement was based on clinical requirements only. Recordings were referenced to a site in white matter. The sEEG signal was sampled at a rate of 1 kHz, filtered between 1 and 250 Hz, and notched at 50 Hz (Neuracle Technology Co., Ltd., Beijing, China). Stimulus-triggered electrical pulses were recorded simultaneously with sEEG signals to precisely synchronize the stimulus with electrophysiological signals.

Patients performed the experiment in a quiet and dim environment. The stimuli were presented on a 24-inch LED screen (Admiral Overseas Corporation, refresh rate of 144 Hz, resolution of 1920 × 1080), and eye position data were obtained by an infrared eye tracker (Jsmz EM2000C, Beijing Jasmine Science & Technology, Co., Ltd.), sampling at 1 kHz. The experimental paradigm was presented by MATLAB (The MathWorks) and Psychtoolbox-3 (PTB-3; Brainard and Pelli). Patients regularly took standard medication for epilepsy treatment during the experimental period.

All patients provided informed consent to participate in this study. The ethics committee of the Chinese PLA General Hospital approved the experimental procedures.

### Electrode localization

Electrode locations were determined by coregistering the postoperative computed tomography (CT) scans with the preoperative T1 MRI using a standardized mutual information method in SPM12 ^40^. Cortical reconstruction and parcellation (with the Desikan-Killiany atlas) were conducted with FreeSurfer ^41^. To calculate the MNI coordinates of recording sites, nonlinear surface registration to the MNI ICBM152 template was performed in SPM12.

### Experimental task

As illustrated in Figure 1, a trial started when a fixation point (0.5° × 0.5°, white cross) appeared at the center of the screen (radius of the eye position check window = 4°, shown as the dotted circle). After the subject fixated on the fixation point for 600 ms, a cue stimulus (Gabor grating, 2 × 2° circle) was presented for 50 ms at a fixed position (7°) on the left (or right) side of the fixation point. The location where the grating appeared (left or right) was opposite to the hemisphere where the electrodes were implanted. There were four patients with electrodes in both hemispheres; the grating was set on the right side for 3 patients and on the left side for one patient. In 70% of the trials, the grating contrast (Weber contrast, see ^36^) was maintained near the subject’s perceptual threshold by a 1 up/1 down staircase method, and the step was 0.39% of the highest contrast of the screen. In 10% of the trials, the stimulus contrast was well above the threshold, and in the other 20% of the trials, the stimulus contrast was 0, namely, no stimulus appeared. After another 600 ms delay, the color of the fixation point turned red or green, and two saccade targets (1 × 1°, white square) appeared at fixed positions (10°) on the left and right sides of the fixation point. If the grating was seen, a green fixation point indicated that participants should make a saccade to the right target, whereas a red fixation point indicated that participants should make a saccade to the left target. If the grating was not seen, the rule of saccadic direction was inverted. Gratings with high contrast (well above the perceptual threshold) and zero contrast (grating absent) served as control conditions to evaluate the understanding and performance of the task. Before data were collected, patients completed 1-2 training sessions, and the contrast perceptual threshold was determined for each subject. During data collection, each session consisted of 180 trials, and the intertrial interval (ITI) was 800 ms.

### Data analysis

Each patient completed 5-7 sessions. We excluded trials in which patients broke fixation during the fixation period (eye position outside of the 4 × 4° check window for more than 100 ms) or the saccadic latency exceeded the maximum response duration (2000 ms for Patients 1-5, 5000 ms for Patients 6-9, according to the patient’s performance) after the target appeared. For all patients, more than 85% of the trials were included in further analysis.

For each session, according to the patients’ reports of being aware or not aware of the presence of gratings, we divided the trials into aware and unaware trials. According to the percentage of patients reporting ‘awareness’, we further divided the grating contrast into 3 levels: high contrast (awareness percentage > 75%), near-threshold contrast (25% ≤ awareness percentage ≤ 75%), and low contrast (awareness percentage < 25%). Due to the small number of low contrast-aware and high contrast-unaware trials, we focused on trials under four awareness conditions: low contrast-unaware (LU), near threshold-unaware (NU), near threshold-aware (NA) and high contrast-aware (HA). On average, there were 228/332/328/147 trials (22.03/32.08/31.69/14.20%, averaged across patients) in the LU/NU/NA/HA conditions, respectively. Since we were interested in the effect of awareness on patients’ behavior and neural activity, we focused on the comparison between NA and NU conditions in the subsequent analysis. The LU and HA conditions were used as the control conditions to verify patients’ understanding of the task.

### Behavioral data analysis

The psychophysical curve was fitted using the following formula:

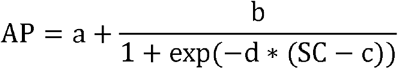

where AP indicates the ratio of reported awareness, SC indicates the stimulus contrast, and a-d indicate individual fitting parameters.

For population psychometric curve fitting, adjacent contrast levels were averaged, resulting in 8 contrast levels in each session.

### LFP data analysis

#### Data preprocessing

Data preprocessing was performed by the software package EEGLAB^42^ in MATLAB. Data were filtered to 1-250 Hz and notch-filtered at 50, 100, 150, 200, and 250 Hz with an FIR filter. Epochs were extracted from -490 ms before to 1299 ms after grating onset. Bad recording sites were discarded from the analysis based on visual inspection of ERP activity and power spectral density (PSD) analysis. Each recording site considered a bipolar reference, i.e., rereferenced to its direct neighbor.

#### Quantitative definition of visual awareness-related activity at recording sites

We first identified visual consciousness-related sites. For all recording sites, we compared the LFP amplitude between the NA and NU conditions in an interval of 0-650 ms after grating onset, in which statistical analysis was conducted for each time point. We defined a recording site as awareness-related if there was a significant difference (p < 0.01, FDR corrected for time points and channels, independent t test) between NA and NU conditions that persisted for more than 20 ms. We also defined the start time of divergence in the LFP between NA and NU conditions as the divergence onset time (DOT).

#### Distribution fitting

The fitting formulas were as follows.

Unimodal distribution:

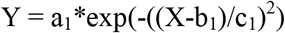

where Y indicates probability density; X indicates DOTs; and a_1_, b_1_, and c_1_ indicate individual fitting parameters.

Bimodal distribution:

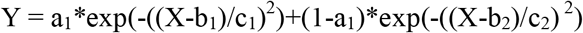

where Y indicates probability density; X indicates DOTs; and a_1_, b_1_,c_1_, b_2_, and c_2_ indicate individual fitting parameters;

#### Population activity calculation

The biological characteristics of LFP polarity are not yet clearly understood^43^. To simplify this complex issue, we defined the change in LFP magnitude during the delay period in our task as reflecting consciousness-related activity, regardless of whether its actual value was positive or negative. Therefore, in the population activity calculation, if the polarity of consciousness-related activity was negative, the time series was multiplied by -1 to unify the polarity across all consciousness-related activity.

#### Brain Region Selection

Due to the sparse sampling of iEEG electrodes in patients, the number of recording sites in different brain regions may vary greatly. To ensure the reliability of the results, we screened the brain regions for inclusion in analysis according to the following criteria:

1. Screening of brain regions to include in the analysis: the brain region had to have ≥10 recording sites in ≥ 3 patients. These regions included the superior frontal gyrus, middle temporal gyrus, rostral middle frontal gyrus, superior parietal gyrus, insula, caudal middle frontal gyrus, caudal anterior cingulate, inferior parietal gyrus, pars opercularis, pars triangularis, thalamus, amygdala, caudate, and white matter.
2. Screening the brain regions to calculate the proportion of early sites: the brain region had to include early sites in ≥ 3 patients.
3. Screening the brain regions to calculate the proportion of late sites: the brain region had to include late sites in ≥3 patients.

### Topographic display of early and late awareness-related activity

To visualize the locations of early and late awareness-related activity, we used the Brainstorm^44^ package of MATLAB to generate topographic maps of the proportions of early and late consciousness-related sites, respectively. The results are shown on the ICBM152 cortical surface. This is a simple 3D interpolation with Shepard’s method (weights decreasing with the square of the distance), in which the vertices of the cortical surface were ignored if they were more than 15 mm from the center of each sEEG contact.

### Lateralized analysis of awareness-related activity

To assess whether awareness-related activity was differentially distributed between the two hemispheres, we calculated the proportion of recording sites with awareness-related activity over all recording sites within each side of the PFC. We repeated this calculation with a 1 ms time bin in the period of 0-650 ms after grating onset.

### Connectivity Analysis

The time-resolved phase-locking value (PLV) was calculated by using the brainstorm toolbox^44^. Time-frequency transformation of the LFP signal was conducted with the Hilbert transform method. The time-resolved PLV reflects whether the phase difference between two LFP signals (i.e., LFP activity in two recording sites) is consistent across trials with respect to the grating onset (i.e., the output is a mean across trials per time point)^45^. PLV values range between 0 (for totally random) and 1 (for perfect phase locking).

The spectral Granger causality was calculated with the brainstorm toolbox^44^. Time-frequency transformation of the LFP signal was conducted with the Hilbert transform method. For individual patients, the average GC of the early-to-late direction was calculated by averaging the GC values of the pairs between early and late activity; the average GC of the late-to-early direction was calculated by averaging the GC values of the pairs between late and early activities.

## Acknowledgments

We thank the patients for volunteering to take part in the study. This study was funded by STI2030-Major Projects+2021ZD0204300. National Natural Science Foundation of China (32030045, 32061143004).

## Data and software availability

Electrophysiological data were analyzed using MATLAB in conjunction with the toolboxes mentioned above. As we have several ongoing studies based on this dataset, the dataset and customized MATLAB analysis scripts are available upon reasonable request from Ming-sha Zhang.

## Author Contribution

Zepeng Fang: conceptualization, methodology, investigation, software, formal analysis, visualization, writing original draft, writing—review and editing, validation, data curation;

Yuanyuan Dang: conceptualization, methodology, investigation, software, formal analysis, visualization, writing review and editing, validation, data curation;

Xiaoli Li: methodology, software, writing review and editing, validation;

Qianchuan Zhao: methodology, writing review and editing;

Mingsha Zhang: conceptualization, resources, writing review and editing, supervision, funding acquisition;

Hulin Zhao: conceptualization, methodology, software, resources, writing review and editing, supervision, project administration.

## Statistical analyses

See in Methods.

